# *Flavoceraceomyces* (NOM. PROV.) (Irpicaceae, Basidiomycota), encompassing *Ceraceomyces serpens* and *Ceriporia sulphuricola*, and a new resupinate species, *F. damiettense*, found on *Phoenix dactylifera* (date palm) trunks in the Nile Delta of Egypt

**DOI:** 10.1101/2020.07.16.206029

**Authors:** Hoda M. El-Gharabawy, Caio A. Leal-Dutra, Gareth W. Griffith

## Abstract

The taxonomy of Polyporales is complicated by the variability in key morphological characters across families and genera, now being gradually resolved through molecular phylogenetic analyses. Here a new resupinate species, *Flavoceraceomyces damiettense* (NOM. PROV.) found on the decayed trunks of date palm (*Phoenix dactylifera*) trees in the fruit orchards of the Nile Delta region of Egypt is reported. Multigene phylogenetic analyses based on ITS, LSU, EF1α, RPB1 and RPB2 loci place this species in Irpicaceae, and forming a distinct clade with *Ceraceomyces serpens* and *Ceriporia sulphuricolor*, which we also incorporate into a new genus *Flavoceraceomyces* (NOM. PROV.). The honey-yellow basidiomes with white margins and presence of crystal-encrusted hyphae in the hymenium and subiculum are distinctive features of *Flavoceraceomyces* (NOM. PROV.), despite variability in hymenium morphology and presence of clamp connections and cystidia, as noted for other genera within Irpicacae. *F. damiettense* is hitherto consistently associated with date palms killed by the red palm weevil *Rhynchophorus ferrugineus*, a highly damaging and invasive pest, recently spread to the Mediterranean region. *F. damiettense* causes rapid wood decay by a potentially unusual white-rot mechanism and may play a role in the damage caused by *R. ferrugineus*.

## Introduction

The taxonomic placement of many taxa within Polyporales has undergone substantial revision in light of molecular phylogenetic evidence, coordinated through focused initiatives such as the PolyPEET project (https://wordpress.clarku.edu/polypeet/) (Binder et al. 2013; Floudas and Hibbett 2015; Justo et al. 2017). It is by now clear that resupinate or corticioid fungi can be difficult to classify even to family level based on morphological characteristics. However, multigene phylogenies now provide a robust backbone at family level but there is an urgent to reclassify many taxa which have been attributed to incorrect genera based on morphological data.

This problem is well-illustrated by the family Irpicaceae Spirin & Zmitr. 2003 (Spirin 2003), a well-supported clade which currently comprises 12 genera (*Byssomerulius, Ceriporia, Cytidiella, Efibula, Emmia, Flavodon, Gloeoporus, Hydnopolyporus, Irpex, Leptoporus, Meruliopsis, Trametopsis*) (Justo et al. 2017). Within multigene phylogenies of Irpicaceae, it is also apparent that several clearly delineated clades within this family remain to be named (Justo et al. 2017).

The family Irpicaceae also illustrates the fundamental problems associated with the classification of Polyporales based on morphological traits. In terms of macromorphology, three forms of basidiomes (pileate [*Trametopsis*], resupinate [*Gloeoporus*] and stipitate [*Hydnopolyporus fimbriatus*]) and four states for hymenophore configuration (poroid, daedaleoid/lamellate, smooth, hydnoid) are found (Justo et al. 2017; Sjökvist et al. 2012). A similar variation is evident in the diversity of hyphal systems (mostly monomitic but some dimitic [*Flavodon*/*Irpex*/*Trametopsis*]), which affects the consistency and longevity of fruiting bodies. Decay mode of Polyporales, is one of the most stable characters that has been used as the basis for segregating genera (Gilbertson and Ryvarden 1986; Ryvarden 1991) but within Irpicaceae, whilst most are white rotting, a single genus [*Leptoporus*] exhibits brown rot decay (Justo et al. 2017). With regard to other microscopic traits, most members of the family are rather nondescript, with consistently smooth hyaline spores but variation bother within and between genera with regard to the presence of cystidia and clamped septa (e.g. cystidia in *Irpex, Emmia* and others; clamp-connections in *Gloeoporus* and others).

These various morphological transitions have occurred repeatedly within this lineage (Floudas and Hibbett 2015; Miettinen et al. 2016), making the construction of any dichotomous key based on morphological characteristics very unwieldy. Recognizing smaller, well-supported clades as independent families would not result in a more straightforward morphological grouping of these taxa.

In this paper we describe a new resupinate fungus found on decaying trunks of *Phoenix dactylifera* trees killed by red palm weevil *(Rhynchophorus ferrugineus*). Based on the genetic and morphological similarities of this new species to two other ‘orphan’ resupinate species (*Ceriporia sulphuricolor, Ceraceomyces serpens*), we include all three taxa in a new genus which we name *Flavoceraceomyces* (NOM. PROV.).

## Methods

### 1. Sampling and Morphological studies

Basidiome samples were collected during a survey for wood-inhabiting fungi across orchards, and gardens of the North Nile Delta region of Egypt (2013-2020). Isolation was conducted from basidiome tissues (Stalpers 1978) at the Microbiological laboratory of Faculty of Science, Damietta University. Pure cultures were obtained on Potato dextrose agar (PDA) and 3% Malt extract agar (MEA), routinely incubated at 28°C. Cultures were stored at 4°C, on agar slopes and frozen at −80°C in 10% glycerol. Radial growth rate was quantified on MEA in 90 mm petri dishes, with mycelial plugs of actively growing cultures placed at the edge of the dish, according to the method of (Adaskaveg and Gilbertson 1986). Optimal growth temperature was investigated across a range of temperatures (20-41°C). Culture compatibility tests were carried out for different isolates on MEA at 30°C for 3 weeks (Worrall 1997). Vouchers from the samples were deposited at Aberystwyth University Herbarium (ABS). Herbarium acronyms follow Index Herbariorum (http://sweetgum.nybg.org/science/ih/).

The basidiome surface was observed with a dissecting microscope (Prior model 29362) at 50x. Basidiome sections were investigated by light microscopy (Olympus BX51M) mounted in 5% KOH, cotton blue or Melzer’s reagent at 1000x magnification. Photomicrographs recorded with a Nikon Coolpix 995 digital camera. Measurements were taken using an objective micrometer or calibrated ocular.

Colony characters as colour, shape and size of hyphae and type of septa were checked after 1,2,4 weeks of incubation on MEA plates at 28°C (Stalpers 1978). Hyphae were mixed with 20 μl of Calcofluor solution (200 μg/ml [w/v] in distilled water to stain the chitin cell walls), then visualized by epifluorescence microscopy (Olympus BX51). The spore shape index (Q; length/diameter) was calculated for 10 spores (Wu 1990). Scanning electron microscopy was performed at IBERS, Aberystwyth University. Sections of air-dried samples were mounted directly on the surface of carbon stubs and coated with platinum and palladium (Pt/Pd; 4 nm thick layer) mixture using a High Resolution Sputter Coater (Agar Scientific Ltd, UK) at 20 mA under vacuum, and the thickness was monitored with a quartz crystal micro-balance thickness controller. SEM was undertaken with a Hitachi S-4700 field emission scanning electron microscope (Hitachi, Tokyo, Japan) with the following emission settings: 10 μA/1500V and using Mixed (M) or upper (U) detectors.

### 2. DNA extraction, PCR amplification and sequencing

Genomic DNA was extracted from pure cultures using a CTAB-based method (Griffith and Shaw 1998). PCR amplifications of several parts of DNA were carried out using different primer pairs. PCR reactions were performed in a volume of 25 μl (13 μl of Dream Taq [Green PCR Master mix] and 2 μl (ca. 1-10 ng) genomic DNA) using a thermal cycler (PTC-100 Techne, UK). PCR cycling conditions were as follows: one cycle of initial denaturation stage at 96°C for 5 min, 35 amplification cycles [Denaturation at 95°C for 30 sec, annealing at 53°-58°C (see below) for 30 sec, extension at 72°C for 45 sec], then final extension at 72°C for 5 min. The following primer pairs were used for different loci: **ITS1, 5**.**8S, ITS2** (55°C anneal) (ITS1F [CTT GGT CAT TTA GAG GAA GTA A] and ITS4 [TTC TTC GCT TAT TGA TAT GC]) (Schoch et al. 2012); **ITS2 and LSU (D1/D2 region** (53°C anneal)(ITS3 [GCA TCG ATG AAG AAC GCA] / HyglonR1 [TAA AGC CAT TAT GCC AGC ATC]) (Detheridge et al. 2016); **RPB1** (55°C anneal) (rpb1:aAf [GAG TGT CCG GGG CAT TTY GG] / rpb1:i2.2f [CGT TTT CGR TCG CTT GAT] and rpb1:aCr [ARA ART CBA CHC GYT TBC CCA T] / rpb1:940R [CTT CGT CYT TCG AAC GYT TRT A]) (Binder et al. 2010); **EF1α** (58°C anneal) (EF1-1018F [GCY CCY GGH CAY CGT GAY TTY AT] / EF1-1620R [ACH GTR CCR ATA CCA CCR ATC TT]) (Rehner and Buckley 2005).

PCR-DNA products were purified using spin column PCR purification kit (NBS Biologicals Ltd., Huntingdon, UK) and visualized on 1.5% agarose gel by gel electrophoresis system. The samples were then sent for Sanger sequencing at the IBERS Translational Genomics Facility (Aberystwyth University).

DNA sequences and chromatograms were curated and assembled using Geneious R10 (Kearse et al. 2012). The sequences generated in this study have been submitted to GenBank (Table 1).

**Table 1.**
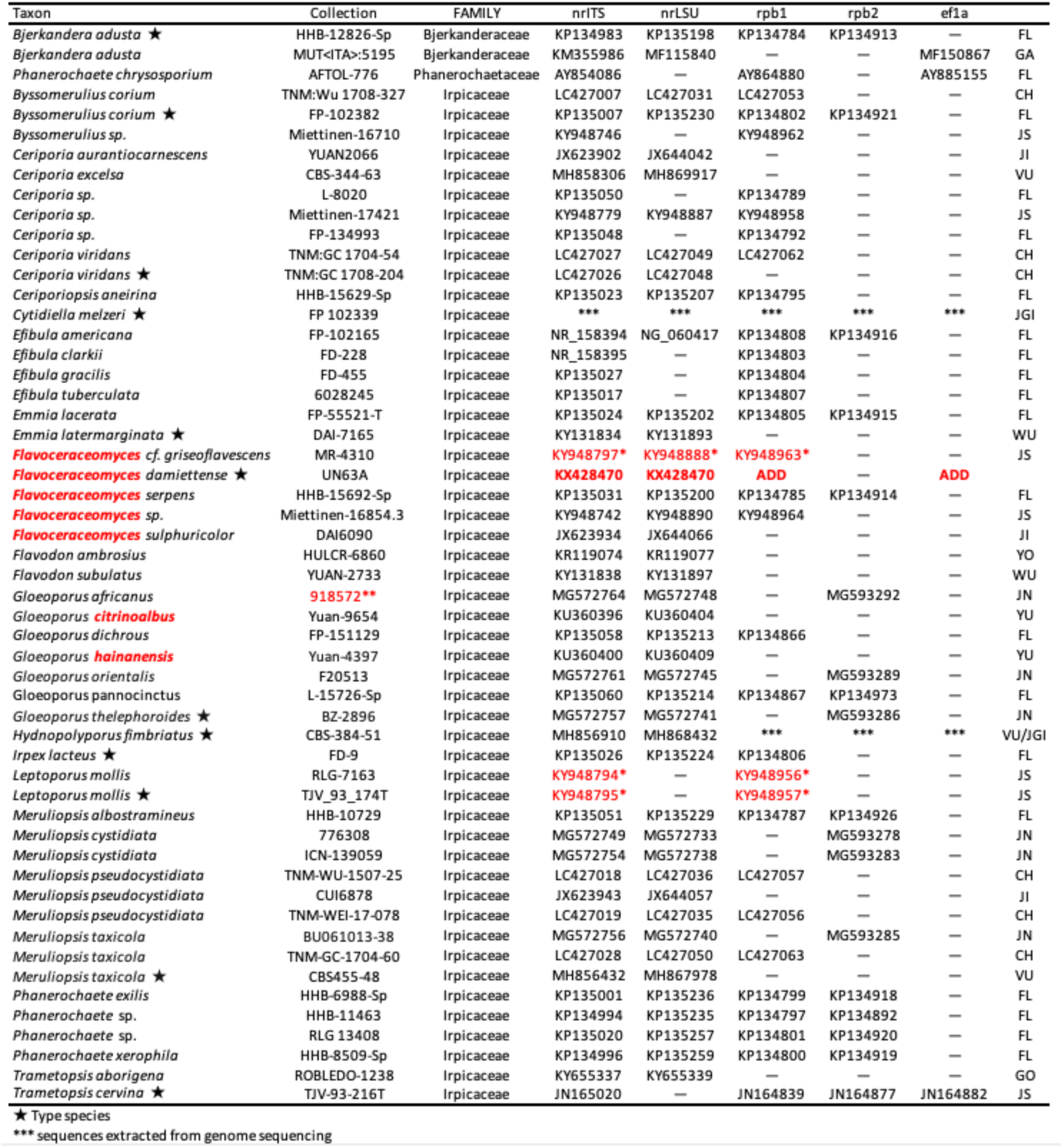
Details of sequences used in the phylogenetic analyses. Sequence data generated in the present study is indicated in bold font. Sequence sources are indicated as follows: CH (Chen et al. 2020); FL (Floudas and Hibbett 2015); GA (Garzoli et al., unpublished); GO (Gómez-Montoya et al. 2017); JGI (Joint Genome Institute); JI (Jia et al. 2014); JN (Jung et al. 2018); JS (Justo and Hibbett 2011); VU (Vu et al. 2019); WU (Wu et al. 2017); YO (Li et al. 2015); YU (Yuan et al. 2016).

In addition to sequences submitted to GenBank from previous phylogenetic studies of Irpicaceae (Table 1), sequences from *Cytidiella melzeri* (FP 102339) and *Hydnopolyporus fimbritatus* (CBS 384.51) were extracted from next-generation sequencing data deposited on NCBI database (SRX3006953-SRX3006967 and SRX2124808-SRX2124809, respectively) using an in-house developed pipeline.

### 3. Phylogenetic analyses

A dataset containing 169 sequences from 54 samples including members of Irpicaceae and three Phanerochaetaceae as outgroup was created including ITS (54 sequences), LSU (54 sequences), EF1α (6 sequences), RPB1 (35 sequences), and RPB2 (20 sequences). GenBank accession numbers and voucher details for the sequences used in these analyses are provided in Table 1.

For all the following analyses the ITS region was divided into three subsets ITS1, 5.8S and ITS2. Only the CDS from the protein-coding genes were used except for the third intron from RPB1 that was included in a separate subset. Thus, eight subsets were created: ITS1, 5.8S, ITS2, LSU, EF1α, RPB1, RPB2 and RPB1_I3. Each subset was aligned separately with MAFFT v7.311 (Katoh and Standley 2013) using the E-INS-i algorithm for ITS1, ITS2 and RPB1_I3, and L-INS-i for the remaining subsets. The alignments were curated manually with AliView v1.5 (Larsson 2014) and trimmed to remove uneven ends. With ITS1, ITS2 and RPB1_I3 alignments, morphological matrices were constructed coding the indels as morphological characters using the simple indel coding (Simmons and Ochoterena 2000) implemented in SeqState (Müller 2005) and the nucleotide alignments were then trimmed with trimAl v1.4.rev22 (Capella-Gutiérrez et al. 2009) with the option -gappyout to remove unalignable regions. Moreover, the protein-coding sequences were analysed in three partitions accounting for each codon position. Therefore, 17 partitions were used for the phylogenetic estimations: five nucleotide partitions (5.8S, LSU, ITS1, ITS2, RPB1_I3), three morphological (ITS1_indel, ITS2_indel, RPB1_I3_indel) and nine accounting for each codon position (EF1α_1, RPB1_1, RPB2_1, EF1α_2, RPB1_2, RPB2_2, EF1α_3, RPB1_3, RPB2_3).

Maximum-likelihood tree reconstruction was performed with IQ-TREE v2 (Minh et al. 2020). The best-fit evolutionary models and partitioning scheme for this analysis were estimated by the built-in ModelFinder (option -m MF+MERGE) allowing the partitions to share the same set of branch lengths but with their own evolution rate (−p option) (Chernomor et al. 2016; Kalyaanamoorthy et al. 2017). Branch supports were assessed with ultrafast bootstrap (UFBoot; 1000 replicates) (Hoang et al. 2018) allowing resampling partitions and then sites within these partitions to reduce the likelihood of false positives on branch support (option --sampling GENESITE) (Gadagkar et al. 2005). Additionally, ultrafast jackknife (1000), SH-aLRT test (1000) (Guindon et al. 2010) and non-parametric bootstrap (100) were performed.

Bayesian Inference (BI) was conducted with MRBAYES v3.2 (Ronquist et al. 2012) with two independent Markov Chain Monte Carlo (MCMC) runs, each one with four chains and starting from random trees. The best-fit evolutionary models and partitioning scheme for these analyses were estimated as for the ML analysis but restricting the search to models implemented on MRBAYES (options -m TESTMERGEONLY -mset mrbayes). Chains were run for 10 million generations with tree sampling every 1000 generations. The burn-in was set to 25% and the remaining trees were used to calculate a 50% majority consensus tree and Bayesian Posterior Probability (BPP). The convergence of the runs was assessed on TRACER v1.7 (Rambaut et al. 2018) to ensure the potential scale reduction factors (PSRF) neared 1.0 and the effective sample size values (ESS) were sufficiently large (> 200).

Nodes with BPP ≥0.95 and/or UFBoot ≥95 were considered strongly supported. Alignment and phylogenetic trees are deposited in TreeBASE (ID: 26614).

## Results

### 1. Field investigation and sample collection

A resupinate fungus with honey-yellow basidiomes and white margins was initially collected as a single infection on the dead stump of a date palm tree (*Phoenix dactylifera*) in February, 2014 in a fruit farm at Baltim city, Kafr El-Sheikh (31.5764°N, 31.0796°E; a governorate of Nile Delta of Egypt). In more detailed surveys, basidiomes were found to be more widespread and present on dead date palm stumps and logs in several adjacent orchards (2017; five infected stumps in area of 200-250 m^2^). In 2018-2020, basidiomes were also found in orchards 60 km to the east in El-Sinaniah, Damietta (31.4429°N, 31.7798°E), always on dead date palm stumps or fallen trunks (10 different trees across an area of about ca. 400 m^2^). In all cases, the date palm stumps were in dense orchard with dense cover of planted trees (mainly mango, lemon, and date palm). Basidiomes were present throughout the year (Winter temp 10-20°C [RH 60-80%], Summer temp 25-35°C [RH 40-45%]).

Some of the basidiomes were formed on the bark (Fig. 1A/D) and others on decorticated areas of exposed wood (Fig. 1C/E/F) and the basidiomes were associated with advanced white-rot decay. The cause of death of the host trees was *Rhynchophorus ferrugineus* (red palm weevil), whose boreholes were readily visible in some stumps (Fig. 1B/C; arrowed).

**Fig. 1.**
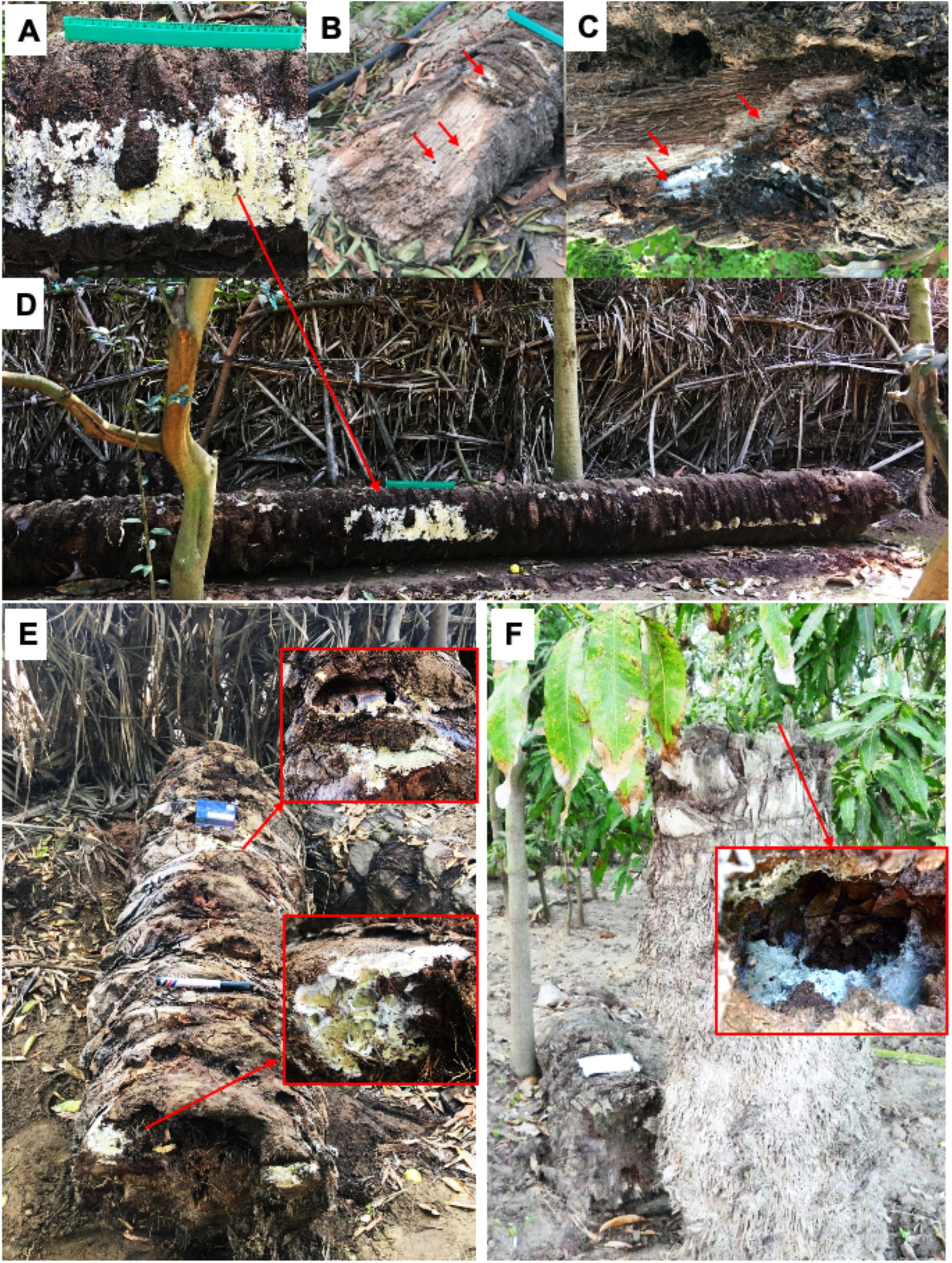
*Flavoceraceomyces damiettense* (NOM. PROV.) infecting date palm trunks. Honey-yellow basidiomes with white margins are visible emerging from the bark (A), often associated with bore holes of red weevil (B, arrowed) and batches of white rot decay (C). Basidiome sometimes formed in long decay columns, also at trunk ends (D) and inside hollowed dead stumps (F).

### 2. Basidiome morphology

The resupinate basidiomes consist of a thin layer of patchy, effuse-resupinate, annual basidiomes, tightly attached to the host, when young and fresh appear as lemon to olive/honey-yellow coloured with a crenulate/papillate/tuberculate hymenium surface, up to 20 × 45 cm in extent, up to 0.3 mm thick (Fig. 2). Sometimes, these appear as a thicker velvety mat, deep yellow coloured, warty (bullate or phlebioid) irregularly crenulated surface with waxy (ceraceous) appearance. Basidiomes have white, thin, loose cobweb-like fibrous margins. With age, the flesh of the basidiome becomes thinner and more compressed, with a pinky-buff or ochraceous to pale brown or olive brown coloured, smoother straight surface, and dull (not waxy) appearance. The flesh is yellowish white, soft, loose when fresh, more fragile with small cracks when dry, becoming dark or brownish colour with addition of KOH solution. No pores were observed on the hymenium. Fresh basidiocarps have a strong but pleasant mushroom-like odour. No mycelial cords or rhizomorphs were observed (Fig. 2).

**Fig. 2.**
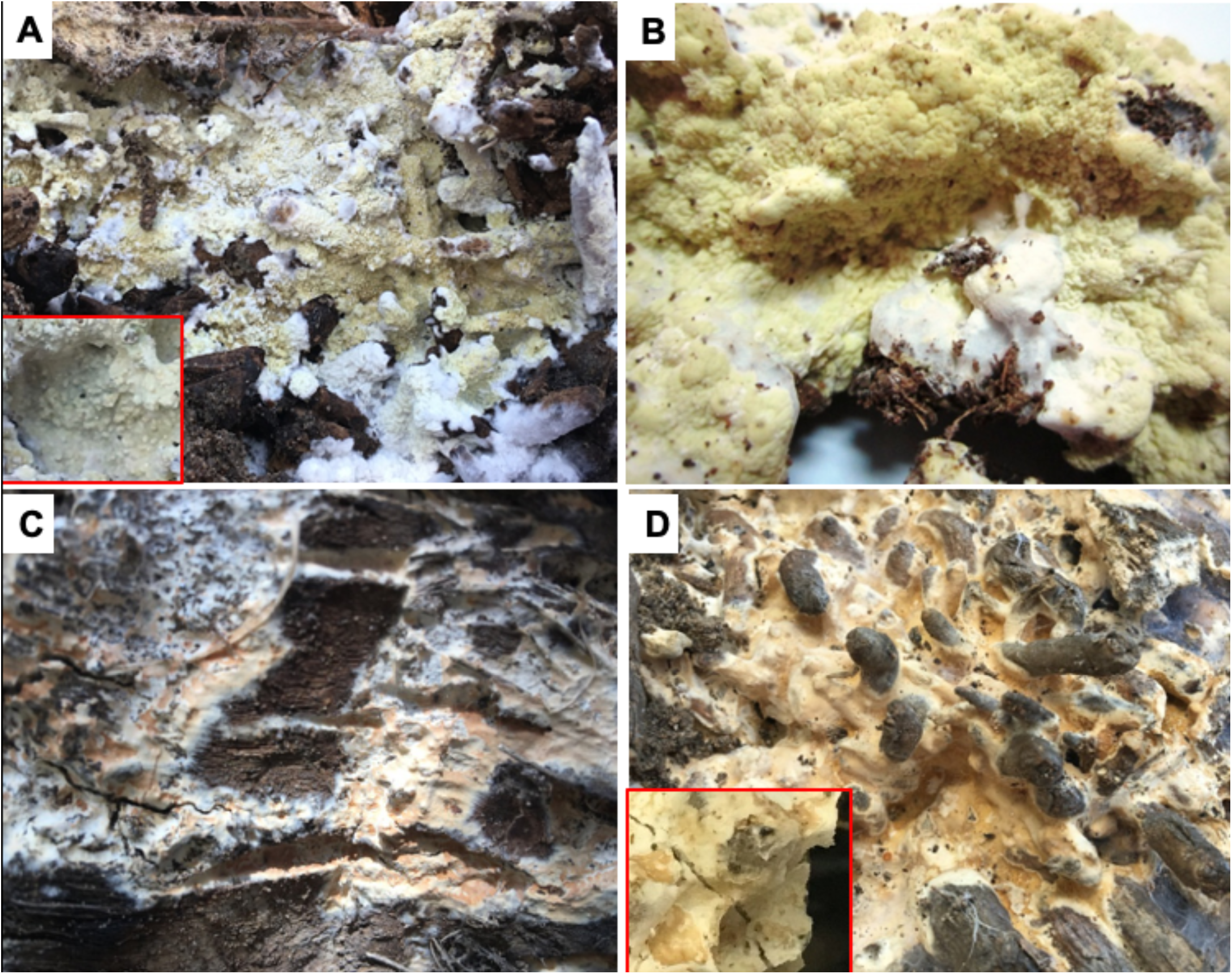
Detailed view of resupinate basidiomes of *Flavoceraceomyces damiettense* (NOM. PROV.) in the field. The resupinate basidiomes are honey yellow coloured with a granulated (papillate) surface and white thin fibrous margin when young (A). Thicker velvety mats with a warty (bullate) surface, irregularly crenulated and a waxy appearance are also found (B). Older basidiomes are buff to pale brown with white margins and cracked (C). The surface become smoother with a dull (not waxy) appearance, with concolorous flesh and soft dense texture (D). Scale bar indicates 1cm.

### 3. Cultural characteristics

Cultures were obtained from eight basidiomes collected from Damietta and Kafr El-Sheikh. In axenic culture (MEA), the mycelium is yellowish or creamy white to pale buff colored, sparse adpressed mycelium, with slow radial growth rate (1.6-2.2 mm/day) (Fig. 3A). On PDA growth is more cottony/downy, with occasionally denser growth on the inoculum plug. Maximal growth rate was obtained at 30°C and no growth was observed above 35°C.

**Fig. 3.**
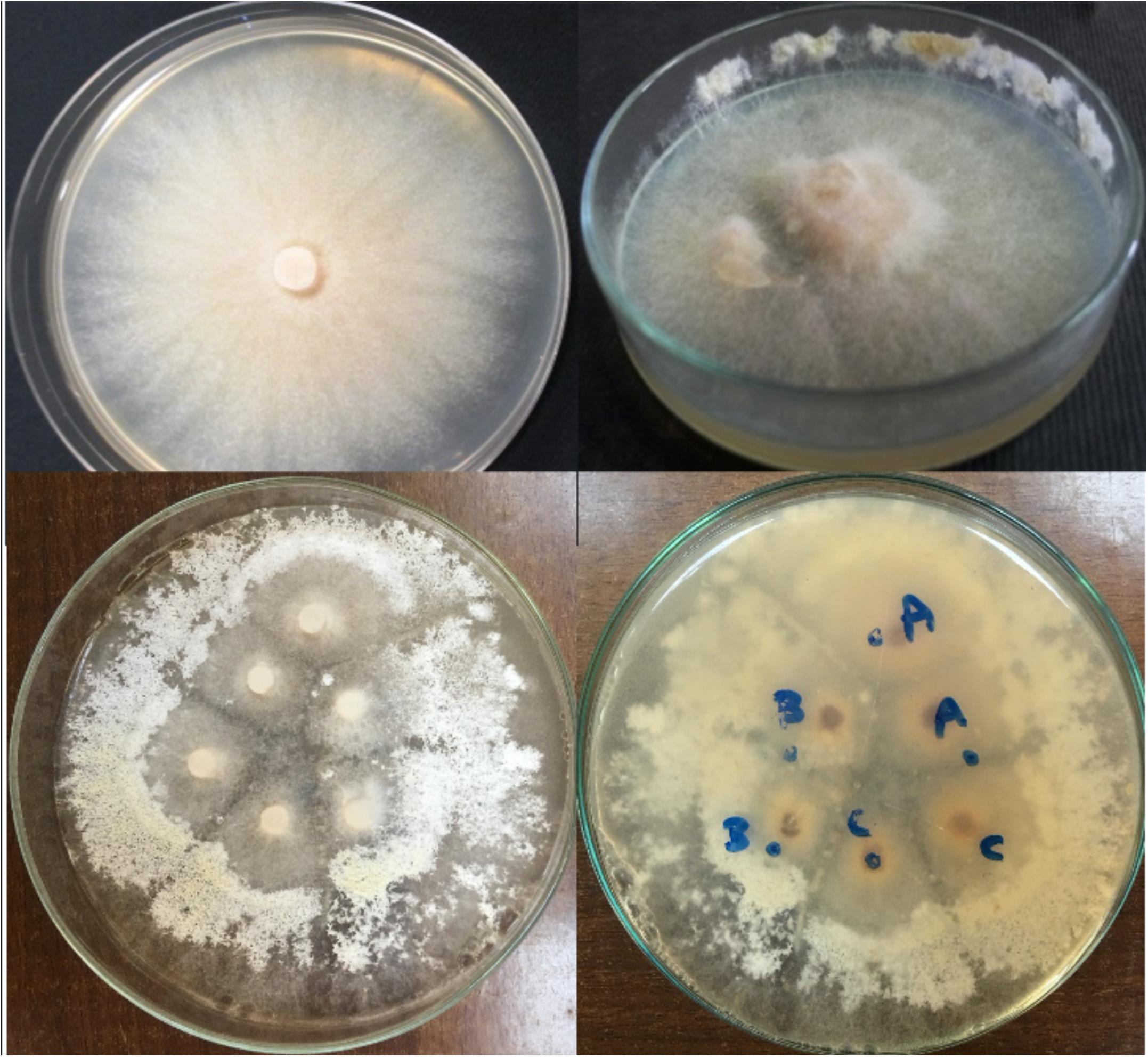
*Flavoceraceomyces damiettense* (NOM. PROV.) in pure culture on PDA. Floccose appearance of young colonies of isolate UN63A (2 weeks old), with slight yellow coloration (A). Colonies incubated in the light exhibited pigmentation similar to that seen in basidiomes (B). Assessment of somatic incompatibility between isolates from Kafr El-Sheikh (UN63A, UN63B, UN63C). Yellow pigmentation after incubation in the light for 4 weeks is visible on the upper surface (C), with zone lines between genetically different cultures (B vs A/C), more clearly visible from beneath (D).

In older cultures (>4 weeks), following growth at ambient light levels, yellowish brown pigmentation developed as a waxy-leathery thin layer at the edges of colonies (Fig. 3B). Thus, there was some differentiation of hymenial structures but basidia and basidiospores were not observed. The pigmentation and differentiation did not occur when cultures were incubated in the dark.

Hyphae from pure cultures are hyaline, thin walled (3-4 μm diameter; Fig 4A/B/C), with large clamps abundant. Chlamydospores, both intercalary and terminal (Fig. 4B/C), were present in older cultures (after 3-4 weeks of growth); these, are numerous, globose or fusiform (7.5-9.5 μm diam) with encrustation by small irregularly shaped crystals visible on some hyphae (Fig. 4D, inset). Amorphous areas of brown resinous material were also visible in some areas but it was not clear whether these were associated with the crystals (Fig. 4D).

**Fig. 4.**
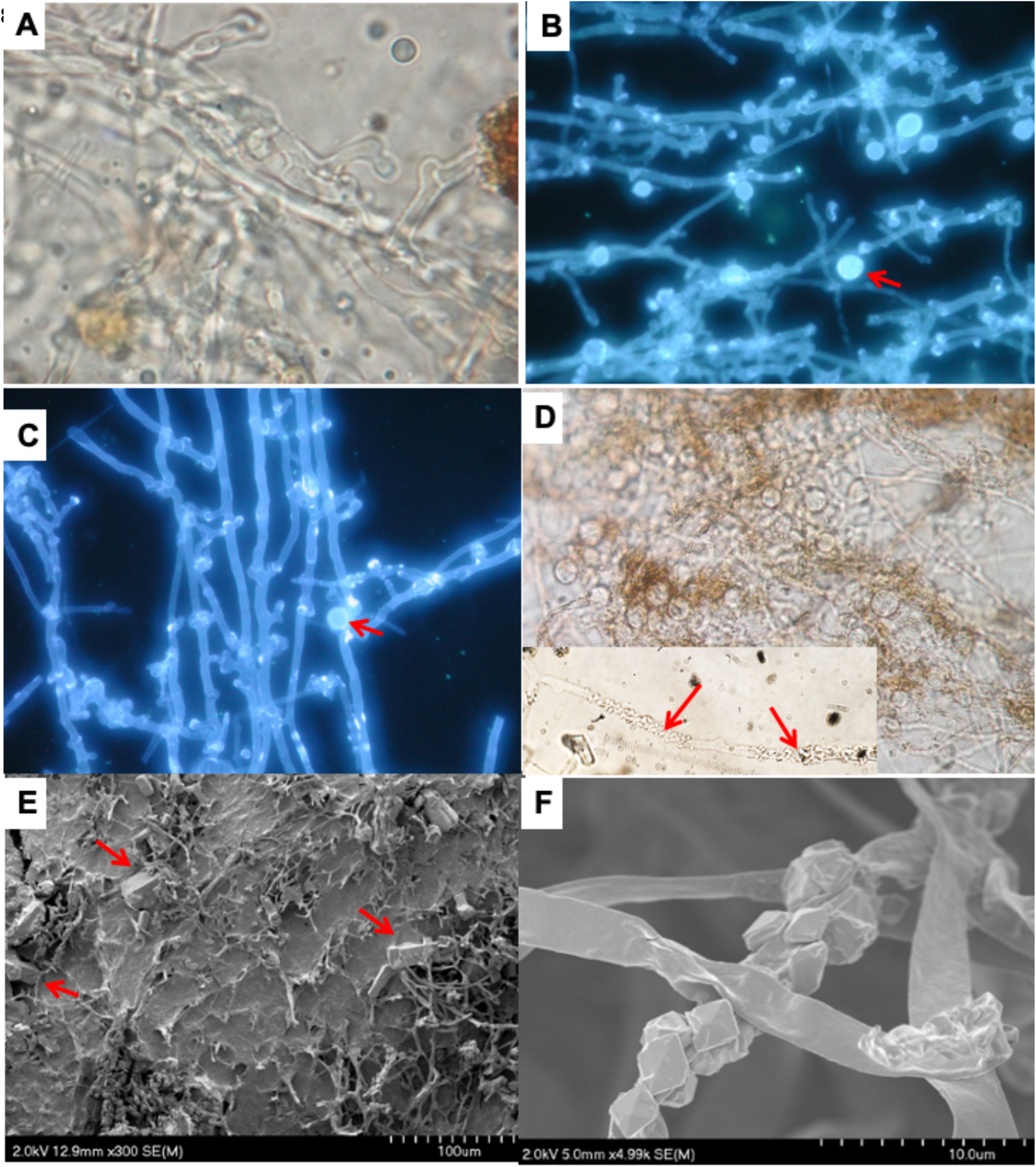
Ultrastructure of *Flavoceraceomyces damiettense* (NOM. PROV.). In pure culture, abundant clamp connections are visible in brightfield (A) and under epifluorescence microscopy with Calcofluor B staining (B,C; arrowed). Spherical-ellipsoid chlamydospores, both intercalary and terminal, are abundant (B-D), alongside brown resinous agglomerations (D) and encrustation by small crystals of individual hyphae (D, inset). In basidiomes examined by scanning electron microscopy, large crystals are visible, embedded in the matrix of the hymenium surface (E; arrowed) and smaller crystals encrusting the surface of hyphae in the subhymenium and subiculum (F). Scale bar indicates 10 μm.

Pairings of isolates obtained from different trees at Kafr El-Sheikh revealed the formation of zone lines due to somatic compatibility (Fig 3. C,D). This result is suggestive of an outcrossing breeding strategy (Griffith and Hedger 1994).

### 4. Basidiome ultrastructure

Basidia, rarely observed, are clavate (6.0-7.5 × 12.0-15.0 μm), smooth, thin-walled, 4-spored (Fig. 5A; SuppFig S1G). Basidioles are similar in shape but smaller (4-5 μm diameter) (Fig. 5A; SuppFig S1G). Basidiospores are short, ovoid to ellipsoid, tear-shaped, smooth, sometimes thick-walled, 3.0-3.5 × 4.0-5.0 μm in size (Q =1.42-1.66), colourless, non-cyanophilous (not staining with Cotton Blue), with no colour change associated with KOH or Melzer’s reagent (i.e. non-amyloid and non-dextrinoid) (Fig. 5A; SuppFig 1H). Cystidia are abundant, long, smooth (not encrusted), hyaline, thin-walled, septate (2-celled), spear-shaped or with a sharply pointed apex (4.0-5.0 × 22-25 μm) (Fig. 5B; SuppFig S1I/J). Cystidioles are fusiform, 3.0-4.0 × 18-22 μm (Fig. 5C; SuppFig S1L).

**Fig. 5.**
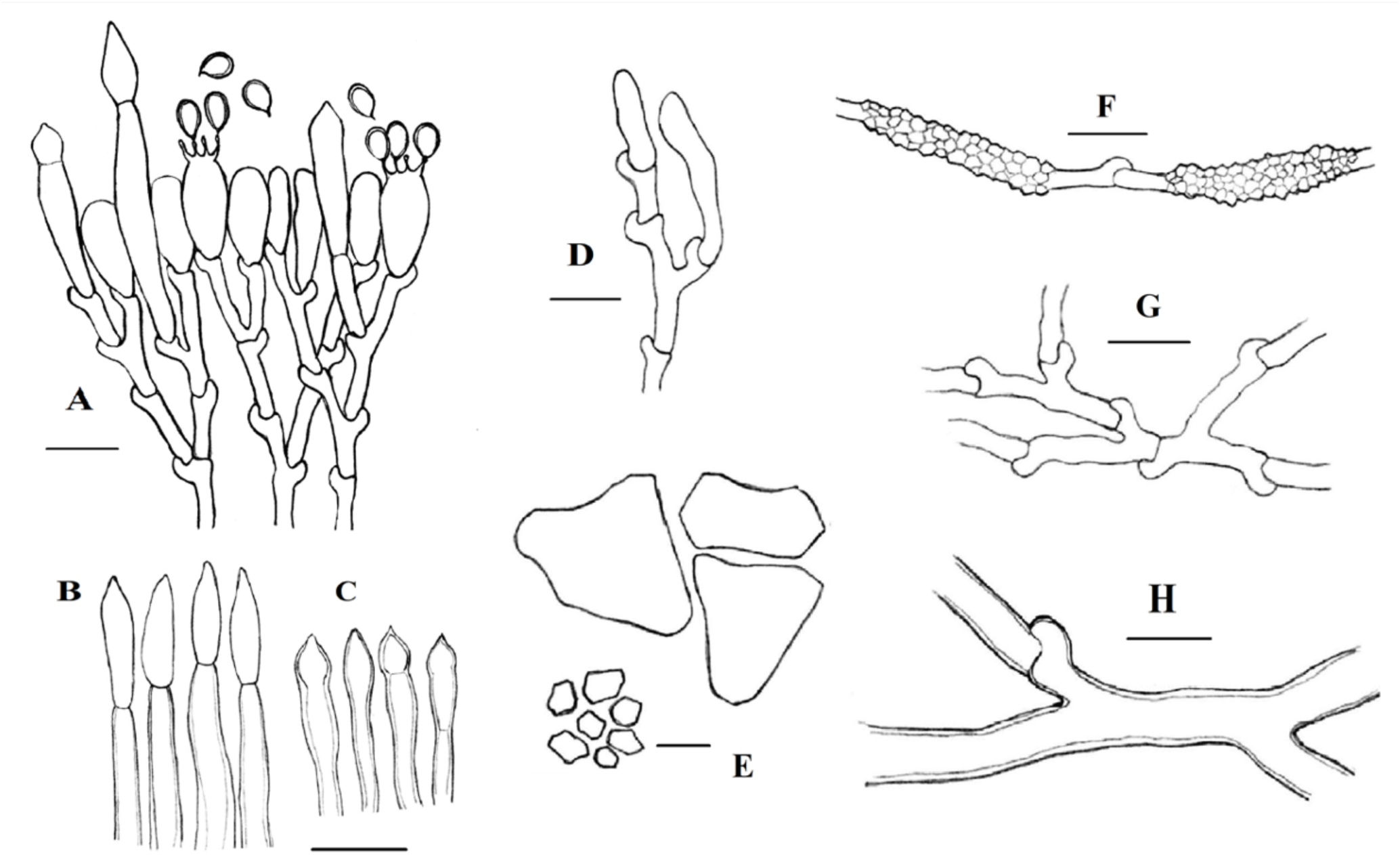
Diagram of microscopic structures of *Flavoceraceomyces damiettense* (NOM. PROV.) basidiome. Hymenium layer with clavate 4-spored basidia, thick-walled basidiospores, smaller basidioles and basal clamps (A). Cystidia are spear-shaped (B) or fusiform (C), with cystidioles also present (D). Large and small hyaline crystals (E) are observed forming encrustation on hyphae (F). Generative hyphae with abundant clamps (G) and pseudo-skeletoid subicular hyphae (H) are present. Scale bar indicates 10 μm.

The subhymenial layer is composed of tightly packed hypha (SuppFig S1E/F), with clamps present on all primary septa, (subicular, subhymenial and sub-basidial). The hyphal system is monomitic; generative hyphae bear abundant clamps (Fig. 5F; SuppFig 1F) and frequent stumpy branches (hyaline and thin-walled, 3.0-4.5 μm diam) (Fig. 5A/B/C). Some generative hyphae are thick-walled to nearly solid (pseudo-skeletoid, sparsely branched, fewer clamps, 6.5-8.0 μm diam) (Fig. 5G).

Large irregularly-shaped hyaline crystals (10-30 × 20-50 μm) are visible on the hymenium surface (Fig. 4E) and also in the subhymenium and subiculum (SuppFig S1C/D/E). More widespread, in the hymenium, subhymenium and subiculum were smaller (1-4 μm) rhomboid-shaped hyaline crystals (Fig. 4F; SuppFig S1F), found mainly forming encrustations of hyphae. The latter were similar to those observed on hyphae in pure cultures. The larger crystals and also the smaller crystals when not forming hyphal encrustations were soluble in KOH but the hyphal encrustations of smaller crystals remained intact in KOH. Brown, resinous agglomerations, similar to those observed in pure cultures, were also visible (SuppFig. S1A) but it was not clear how these were associated with the crystalline deposits. No chlamydospores were seen within basidiome tissues.

**SuppFig. S1.**
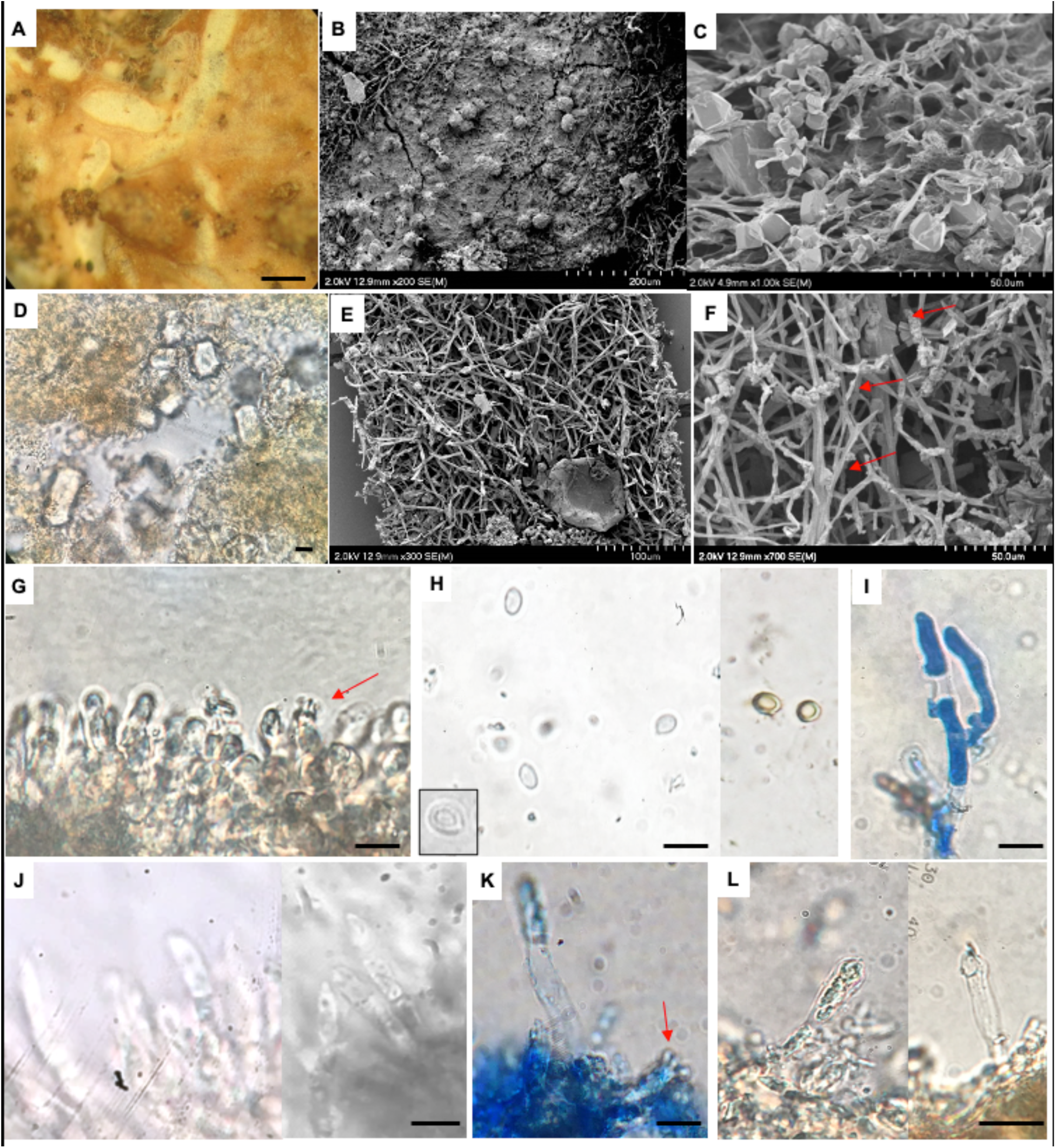
Microscopic features of *Flavoceraceomyces damiettense* (NOM. PROV.) basidiome. A smooth/non-perforated, honey-yellow hymenium encrusted with brown clumps /resinous granules (A: 50x light magnification), which were visible under SEM (B); also clusters of large, amorphous, hyaline crystals observed under SEM (C) and LM (D). In transverse section; the basidiome is composed of a compact mat of mycelia (E) with a monomitic hyphal system consists of three types of generative hyphae, as arrowed (normal generative hyphae with clamps, crystal-encrusted basal hyphae and pseudo-skeletoid hyphae) (F). Clavate 4-spored basidia (arrowed), smaller basidioles (G) are visible on the hymenium, as are sometimes thick-walled ovoid basidiospores (H), showing no reaction with Melzer’s reagent (H; right). Generative hyphae with basal clamps (I; Cotton Blue stain) are visible, as are spear-shaped (2-celled) cystidia (J, K) and fusiform cystidioles (L). Scale bar indicates 10 μm (except A = 400 μm).

### 5. Phylogenetic reconstruction

The final alignment consisted of 54 sequences with 4291 characters and 2133 parsimony-informative sites. The BI analysis converged both runs as indicated by the effective sample sizes (ESS) of all parameters above 4500 and the potential scale reduction factors (PSRF) equal 1.000 for all the parameters according to the 95% highest posterior density interval. Details about evolutionary models and partitioning schemes used and all trees generated in the analyses can be found in SuppData1.

**SuppData1**. Additional details of the phylogenetic analyses, including various trees.

DNA sequence was obtained for the full ITS region of all seven isolates and all were identical (KX428470). Additionally, for isolate UN63A, the D1/D2 region of the large (28S) ribosomal subunit (GenBank KX428470), the EF1α gene (GenBank XXX in progress) and also the RPB1 gene (GenBank YYY in progress) were sequenced. In all cases the sequences obtained were quite distinct from any other sequences present on GenBank but more detailed phylogenetic analysis placed the fungus at the base of the *Byssomerulius* clade, as defined by Floudas and Hibbett (2015) and intermediate between *Meruliopsis* sp. and *Ceriporia* spp. (Fig. 6).

**Fig. 6.**
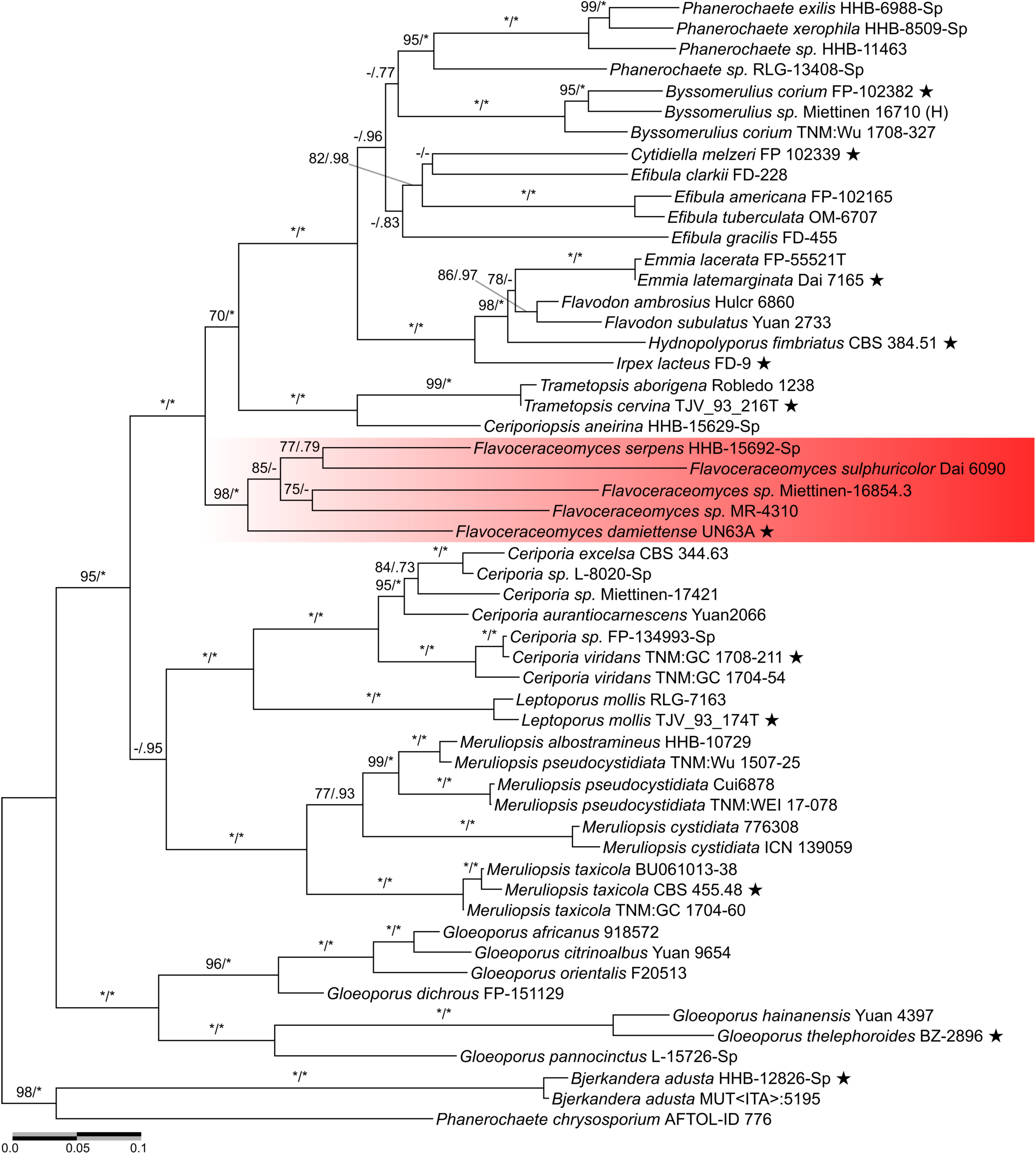
Multigene maximum-likelihood tree of Irpicaceae. Support values are presented as numbers (UFBoot/BPP) on branches and shown only for UFBoot ≥ 70 and BPP ≥ 0.70. Type species for other genera within Irpicaceae are represented by star. Outgroups are *Bjerkandera adusta* and *Terana caerulea* (Phanerochaetaceae). *Flavoceraceomyces* (NOM. PROV.) is boxed in red. Asterisks (*) represent maximum UFBoot/BPP values, dashes (−) represent values below the cut-off threshold (70%), and dots (.) represent ML clades that were not recovered in the BI tree Scale bar indicates number of substitutions per site. More details on supporting values can be found in SuppData1.

Sequences were also extracted from two genome sequence projects (*Cytidiella melzeri* and *Hydnopolyporus fimbriatus*). Alignment and phylogenetic trees are deposited in TreeBASE (ID: 26614).

The clades recovered here within Irpicaceae correspond well with those recovered by Justo et al. (2017)(Fig. 4). Clade /flavoceraceomyces (BS=98; BPP=1) is strongly supported in our phylogeny and occupies a similar position in the phylogeny presented by Justo et al. (2017), basal to the subclade containing *Trametopsis, Efibula, Byssomerulius* and *Irpex*. Within clade /flavoceraceomyces, UN63 occupies a basal position, with the other samples forming two distinct pairs. The detailed morphological descriptions of *F. serpens* and *F. sulphuricolor* (Bernicchia and Niemelä 1998; Ginns 1975) suggest that these two clades represent distinct species; however, the type specimens of these have not been subject to genetic analysis.

#### Taxonomy

***Flavoceraceomyces* H**.**M. El-Gharabawy, Leal-Dutra and G**.**W. Griff**., (NOM. PROV.)

IndexFungorum. IF557789

##### Type species

*Flavoceraceomyces damiettense* H.M. El-Gharabawy, Leal-Dutra and G.W. Griff.

##### Etymology

Derived from the Latin words *flavus* = yellow and *ceraceus* = waxy, due to the appearance of the basidiomes.

##### Diagnosis

Differs from other members of family Irpicaceae due to the presence of hyphae encrusted with crystals in the subiculum and yellow colour of the hymenial surface when fresh.

##### Description

Basidiomes resupinate with smooth, tuberculate, papillate, merulioid or poroid hymenophores, yellow (typically honey-yellow) in colour, often waxy with white margin. Hyphae of subiculum and sometimes hymenium/subhymenium encrusted with crystals which may be associated with darker (brown) resinous granules. Hyphal system is monomitic, sometimes with abundant clamp connections present. Basidia cylindric-clavate, basidiospores hyaline, smooth, ellipsoid, and non-amyloid and non-dextrinoid. Found forming white rot on heavily decayed trunks of woody monocot and dicot angiosperm, as well as coniferous trees. Rhizomorphs and mycelial cords usually absent.

##### Notes

*Flavoceraceomyces* is also defined as the least inclusive clade containing the following pairs of sequences: ITS1-2 spacer regions rDNA (KX428470, JX623934); D1–D2 region of 28S rDNA (KX428470, JX644066). Chlamydospores abundant in old agar cultures of *F. damiettense* and sometimes in *F. serpens* (Ginns 1975).

***Flavoceraceomyces damiettense* El-Gharabawy, Leal-Dutra and G**.**W. Griff**., (NOM. PROV.)

IndexFungorum. IF557790

##### Etymology

The species epithet “*damiettense*” refers to Damietta University (North Nile Delta, Egypt), close to the location where the fungus was first discovered.

##### Diagnosis

Basidiome resupinate, honey-yellow, tuberculate to papillate-warty, with waxy texture, white margin. Basidia clavate (6.0-7.5 × 12.0-15.0 μm), smooth, thin-walled, 4-spored. Basidioles similar in shape but smaller (4-5 μm diameter). Basidiospores short, ovoid to ellipsoid, tear-shaped, smooth, sometimes thick-walled (3.0-3.5 × 4.0-5.0 μm), non-amyloid and non-dextrinoid. Cystidia abundant, long, smooth, hyaline, thin-walled, septate (2-celled), spear-shaped (4.0-5.0 × 22-25 μm). Cystidioles are fusiform, 3.0-4.0 × 18-22 μm. Monomitic hyphal system, with generative hyphae bearing abundant clamps and frequent stumpy branches. Brown, resinous agglomerations and large irregularly-shaped hyaline crystals (10-30 × 20-50 μm) are present on the hymenium surface, subhymenium and subiculum. Smaller (1-4 μm) rhomboid-shaped hyaline crystals are present forming encrustations of hyphae. Differs from other members of this genus in having abundant cystidia and cystidioles.

##### Typification

The holotype (Fig. 2, Fig. 5) was collected from: EGYPT. Kafr El-Sheikh, Baltim (31.5764°N, 31.0796°E; North Nile Delta), growing on fallen trunk of date palm (*Phoenix dactylifera*) killed by *Rhynchophorus ferrugineus*, 14 Feb 2014, coll. HM El-Gharabawy, Voucher UN63A is held at <ABS>(Aberystwyth University biorepository). The ex-type culture UN63A is held in the Aberystwyth University fungal culture collection and also at the Faculty of Science at Damietta University. GenBank: KX428470 (ITS1 spacer, 5.8S rRNA gene, ITS2 spacer, D1–D2 28S rRNA).

##### Additional specimens examined

Four additional basidiomes found on trunks and stumps of fallen date palms killed by *Rhynchophorus ferrugineus* in the same orchard at Baltim, Kafr El-Sheikh (31.5764°N, 31.0796°E), Aug-Dec 2017; vouchers UN63B, UN63C, UN63X, UN63Y; coll. HM El-Gharabawy, held at the Faculty of Science at Damietta University. Ten basidiomes, on trunks and stumps of fallen date palms killed by *Rhynchophorus ferrugineus*, at El-Sinaniah, Damietta (31.4429°N, 31.7798°E), Jan 2018-Feb 2020; vouchers UN1-UN10; coll. HM El-Gharabawy, held at the Faculty of Science at Damietta University.

##### Distribution

Nile Delta region of Egypt. Hitherto only found on fallen and heavily-rotted trunks or stumps of *Phoenix dactylifera* (datepalm).

##### Notes

White rot decay mechanism with secretion of Mn-dependent and Mn independent peroxidases, but only low and transient secretion of laccase (El-Gharabawy et al. 2016).

***Flavoceraceomyces serpens* (Tode ex Ginns) El-Gharabawy, Leal-Dutra and G**.**W. Griff**., (NOM. PROV.) IndexFungorum: IF557791

##### Basionym

*Merulius serpens* Tode, Abh. Hallischen Naturf. Ges. 1: 355 (1783)

##### Synonyms

*Byssomerulius ceracellus* (Berk. & M.A. Curtis) Parmasto, Consp. System. Corticiac. (Tartu): 81 (1968)

*Byssomerulius farlowii* (Burt) Parmasto, Eesti NSV Tead. Akad. Toim., Biol. seer 16(4): 384 (1967)

*Byssomerulius serpens* (Tode) Parmasto, Eesti NSV Tead. Akad. Toim., Biol. seer 16(4): 384 (1967)

*Byssomerulius serpens* f. *pallidus* Parmasto, Eesti NSV Tead. Akad. Toim., Biol. seer 16(4): 384 (1967)

*Byssomerulius serpens* var. *stratosus* (Pilát) Parmasto, Eesti NSV Tead. Akad. Toim., Biol. seer 16(4): 384 (1967)

***Ceraceomyces serpens* (Tode) Ginns, Can. J. Bot. 54(1-2): 147 (1976) (Ginns 1975)** (NB. Date on IF/MB is 1976 but date of paper is actually 1975.)

*Ceraceomerulius serpens* (Tode) Ginns, Cortic. N. Eur. (Oslo) 2: 201 (1997)

*Lilaceophlebia serpens* (Tode) Spirin & Zmitr., Nov. sist. Niz. Rast. 37: 180 (2004)

*Merulius ceracellus* Berk. & M.A. Curtis, Grevillea 1(no. 5): 69 (1872)

*Merulius crustosus* (Pers.) Duby, Bot. Gall., Edn 2 (Paris) 2: 796 (1830)

*Merulius farlowii* Burt, Ann. Mo. bot. Gdn 4: 336 (1917)

*Merulius porinoides subsp. serpens* (Tode) Bourdot & Galzin, Hyménomyc. de France (Sceaux): 348 (1928) [1927]

*Merulius serpens* Tode, Abh. Hallischen Naturf. Ges. 1: 355 (1783)

*Merulius serpens* var. *stratosus* (Pilát) Parmasto, Scripta Bot. 2: 212 (1962)

*Merulius stratosus* Pilát, Bull. trimest. Soc. mycol. Fr. 52(3): 322 (1937) [1936]

*Serpula serpens* (Tode) P. Karst., Bidr. Känn. Finl. Nat. Folk 48: 345 (1889)

*Serpula serpens* subsp. *ceracella* (Berk. & M.A. Curtis) P. Karst., Kritisk Öfversigt af Finlands Basidsvampar, Tillägg 3: 21 (1898)

*Serpula serpens* var. *pinicola* P. Karst., Kritisk Öfversigt af Finlands Basidsvampar, Tillägg 3: 21 (1898)

*Sesia ceracella* (Berk. & M.A. Curtis) Kuntze, Revis. gen. pl. (Leipzig) 2: 870 (1891)

*Sesia serpens* (Tode) Kuntze, Revis. gen. pl. (Leipzig) 2: 870 (1891)

*Xylomyzon crustosum* Pers., Mycol. eur. (Erlanga) 2: 34 (1825)

*Xylomyzon serpens* (Tode) Pers., Mycol. eur. (Erlanga) 2: 31 (1825)

##### Description

(Ginns 1975): 147

##### Notes

This widespread and common species in Europe and North America. The species was placed in the genus *Ceraceomyces* by Ginns (1975). Ginns (1975) distinguished this white-rot forming species from its ‘relative’ *C. borealis* on the basis that the latter formed a brown rot. However, he noted that whilst *C. serpens* formed a white rot of angiosperm hosts, on conifers this was less clear (“incipient brown rot”). The type species of *Ceraceomyces* is *C. tessulatus*, only very distantly related (Amylocorticiales) to *C*.*serpens* which Ginns and other consistently placed intermediate between *Meruliopsis* and *Byssomerulius*. Ginns (1975) also noted that the subhymenial area of basidiomes was “often heavily impregnated with fine granules or a resin-like substance”.

***Flavoceraceomyces sulphuricolor* (Bernicchia & Niemelä) El-Gharabawy, Leal-Dutra and G**.**W. Griff**., (NOM. PROV.)

IndexFungorum. IF557792

##### Basionym

*Ceriporia sulphuricolor* Bernicchia & Niemelä, Folia cryptog. Estonica 33: 15 (1998)

##### Synonym

*Gloeoporus sulphuricolor* (Bernicchia & Niemelä) Zmitr. & Spirin, in Zmitrovich, Malysheva & Spirin, Mycena 6: 36 (2006)

##### Description

(Bernicchia and Niemelä 1998) : 15

##### Notes

This rare species was described from the Adriatic coast of Italy growing on heavily decayed and wet wood inside the trunk of a dead pine, oak or juniper tree (Bernicchia and Niemelä 1998). The type specimen at HUBO was recently transferred to Oslo (O) and isotype is lodged at Helsinki (H). There are hitherto no DNA sequences from the type specimen but more recently a morphologically very similar voucher (Dai 6090), collected on rotten angiosperm wood from Huangshan Mountain, Anhui Province, China (Cui and Jia 2011) was subject to DNA barcoding (ITS and LSU regions of rRNA) by Jia et al. (2014) who reported it to be basal to the main *Ceriporia* clade, with Chen et al. (2020) placing it close to *Ceraceomyces serpens* in their analyses. No cystidia were observed but hyphoid cystidioles (hyphidia) were reported (Bernicchia and Niemelä 1998) but not (Cui and Jia 2011). The hyphae of the subiculum are heavily encrusted, with small (2-4 μm diameter) hyaline crystals (Bernicchia and Niemelä 1998; Cui and Jia 2011). Cui et al. (2011) noted the similarity of voucher Dai 6090 to *Ceriporia subspissa*, collected in Guyana (Aime et al. 2007) but the latter species is reported to bear cystidia in the hymenium and have a deep reddish brown hymenium/subiculum.

## Discussion

Resupinate fungi are among the most important wood-decay fungi. Not only do they contribute to nutrient cycling by decomposing wood debris, but they are also a potentially valuable source of novel pharmacologically-active biomolecules (Stošić-Grujičić et al. 2011; Zapora et al. 2016). During a survey of basidiomycetes growing on North Nile Delta of Egypt, resupinate basidiomes with unusual morphology were found growing on fallen, dead trunks of *P. dactylifera* in a fruit farm.

Following initial DNA barcoding, generic placement of this taxon could not be determined readily from its morphological features because it possessed characters assignable to several genera. However, following more detailed phylogenetic analysis using RPB1 and EF1α locus, UN63A was placed close to *Ceraceomyces serpens* and *Ceriporia sulphuricolor* with strong statistical support. Previous phylogenetic reconstructions including these two species as also placed them in the same clade (Chen et al. 2020; Justo et al. 2017), and it was already apparent that the generic names attributed to these species was incorrect, since the type species of *Ceraceomyces* is in Amylocorticiales, whilst *Ceriporia viridans* (type species for this genus, also in Irpicaceae) is placed close to *Meruliopsis*. It should be noted that there is as yet no DNA sequence data obtained for the type specimen of *C. sulphuricolor* (Bernicchia 6591, from Italy; (Bernicchia and Niemelä 1998)), with our phylogenetic analyses based on voucher Dai6090 from China (Cui and Jia 2011; Chen et al. 2020). However, the morphological descriptions of the two specimens are very similar (Bernicchia and Niemelä 1998; Cui and Jia 2011), except for the absence of hyphoid cystidia and presence of interwoven (rather than parallel) tramal hyphae in voucher Dai6090.

As has been found to be the case with other genera within Irpicaceae and other families within Polyporales (Floudas and Hibbett 2015; Justo et al. 2017), morphological comparisons of the three species which we now place within *Flavoceraceomyces* (NOM. PROV.), revealed relatively few consistent features but all three form yellow basidiomes with a white margin and crystalline encrustations of the hymenium, subhymenium and subiculum are present. However, other features such as presence of cystidia and clamp connections, as well as the macromorphology of the hymenophore surface are not conserved within the genus.

Ginns (1975)(p. 147) observed that *C. serpens* exhibited some unusual traits in wood decay, “with a white rot of angiosperms and what, visually, appears to be an indistinctive white or incipient brown rot of conifers”. In pure culture he observed it to oxidise gallic acid strongly but did not detect laccase activity using gum guaiac substrate. We have previously examined the ligninolytic capability UN63A using the model substrate ABTS (2,2’-azino-bis(3-ethylbenzothiazoline-6-sulfonic acid)) by El-Gharabawy et al. (2016). It was found to secrete Mn-dependent and Mn-independent peroxidase, as well as laccase. Despite its slow growth rate and low enzymatic activity relative to the other Polyporales with which it was compared, it caused rapid loss of substrate dry weight in ash sawdust culture. In nature, *F. damiettense* causes a white-rot on decaying datepalm trunks but enzymatic evidence suggests that it may have an unusual mechanism of wood decay. *C. sulphuricolor* is reported to produce a white rot but to our knowledge no pure cultures of this species exist. However, a culture derived from voucher MR-4310 (*Phlebia* cf. *griseoflavescens*) is available at CFRM.

There have been very few previous studies of macrofungi associated with datepalm logs. Of these, Rattan et al. (1980) was the most detailed with regard to resupinate fungi but none of the organisms they describe (from Iraq) matches the features of *Flavoceraceomyces damiettense* (NOM. PROV.). Egypt is globally the largest grower of date palm and the arrival of red palm weevil in 1993 has caused a drop in production exceeding 30% (Al-Dosary et al. 2016). It is possible that this fungus has only recently arrived in the Mediterranean region, following the spread of the red palm weevil *Rhynchophorus ferrugineus* (since the 1980’s). This highly invasive pest attacks more than 40 palm species and has spread from Asia widely across the Mediterranean region in recent decades (Al-Ayedh 2008; Al-Dosary et al. 2016).

The consistent association of F. damiettense with *R. ferrugineus*. suggests that a mutualistic interaction between these organisms, as has been found in other wood-boring insects (Geib et al. 2008; Kasson et al. 2016; Skelton et al. 2019a; Son et al. 2011) and that the presence of *F. damiettense* (NOM. PROV.) exacerbates the damaging effects of the weevil. For example, the white rot fungus *Donkioporia expansa* (Polyporaceae) is found in consistent association with the deathwatch beetle (*Xestobium rufovillosum*) (Belmain et al. 2002; Campbell and Bryant 1940). Similarly, *Flavodon ambrosius* (also Irpicaceae) is found as mycosymbiont of *Ambrosiodmus* ambrosia beetles (Li et al. 2015), being transported by their vector in specialised mycangia (Simmons et al. 2016) and suppressing the activity of other wood decay fungi (Skelton et al. 2019b). There is hitherto no direct evidence at present that *F. damiettense* (NOM. PROV.) contributes to the tree damage caused by *R. ferrugineus* (subfamily Dryophthorinae, within the same family [Curculionidae; order Coleoptera] as Ambrosia beetles [subfamilies Platypodinae and Scolytinae] and the bark beetles [subfamily Scolytinae]) but the coincidence in the distribution of the two species raises the possibility that the latter may be a targeted vector of the former (Jacobsen et al. 2017).

## Funding

We thank the Egyptian Government for the funding of a Joint-mission PhD programme (2013-2016) and also a postdoctoral fellowship to visit Aberystwyth in 2020. The Institute of Biological, Environmental, and Rural Sciences receives strategic funding from the BBSRC.

## Competing interests

The authors have declared that no competing interests exist.

## Author contributions

HME - undertook fieldwork and labwork; writing of the paper; CALD - Phylogenetic analyses; writing of the paper; GWG - conceived the paper; analysis of data; drafted the manuscript.

## Acknowledgements

We are grateful to Alan Cookson, IBERS, for assistance with Scanning electron microscopy and also to IBERS HPC and Supercomputing Wales (SCW project, is part-funded by the European Regional Development Fund via Welsh Government) for computing support. The Institute of Biological, Environmental, and Rural Sciences receives strategic funding from the BBSRC. We thank Dr. Amira El-Fallal at the Faculty of Science, Damietta University for his help during forays for fungal samples collection, and Dr. Paul Kirk for taxonomic advice.

## SuppData1

### Parameters for phylogenetic analyses

Partitions file with partition schemes and evolutionary models implemented in the Maximum likelihood analyses (IQTREE partitions file):

~~~
#nexus
begin sets;
charset ITS1_ITS2_I3_RPB1 = Irpicaceae_all.fas: 1-883;
charset 58S_EF1a_2_RPB1_2_RPB2_2 = Irpicaceae_all.fas: 884-1042 1044-3418\3;
charset LSU = Irpicaceae_all.fas: 3419-4291;
charset EF1a_1_RPB1_1_RPB2_1 = Irpicaceae_all.fas: 1043-3418\3;
charset EF1a_3 = Irpicaceae_all.fas: 1045-3418\3;
charset RPB1_3_RPB2_3 = Irpicaceae_all.fas: 1657-3418\3;
charset I3_RPB1_indel = Irpicaceae_all_indel.fas:MORPH, 1-98;
charset ITS2_indel_ITS1_indel = Irpicaceae_all_indel.fas:MORPH, 99-329;
charpartition mymodels =
 TPM2+F+I+G4: ITS1_ITS2_I3_RPB1,
 TPM3+F+I+G4: 58S_EF1a_2_RPB1_2_RPB2_2,
 GTR+F+I+G4: LSU,
 TIM+F+I+G4: EF1a_1_RPB1_1_RPB2_1,
 GTR+F+I+G4: EF1a_3,
 TPM2+F+G4: RPB1_3_RPB2_3,
 MK+FQ+ASC+G4: I3_RPB1_indel,
 MK+FQ+ASC+R2: ITS2_indel_ITS1_indel;
end;
~~~

Partitions file with partition schemes and evolutionary models implemented in the Bayesian Inference analysis:

Models and partition schemes:

~~~
GTR+F+I+G4: ITS1_ITS2_I3_RPB1,
HKY+F+I+G4: 58S_EF1a_2_RPB1_2_RPB2_2,
GTR+F+I+G4: LSU,
GTR+F+I+G4: EF1a_1_RPB1_1_RPB2_1,
GTR+F+I+G4: EF1a_3,
GTR+F+G4: RPB1_3_RPB2_3;
MrBayes partitions file:
#nexus
begin mrbayes;
 execute irpicaceae_mb.nex;
 charset ITS1_ITS2_I3_RPB1 = 1101-1982;
 charset 58S_EF1a_2_RPB1_2_RPB2_2 = 330-488 490-1100\3 2857-4619\3;
 charset LSU = 1983-2855;
 charset EF1a_1_RPB1_1_RPB2_1 = 489-1100\3 2856-4619\3;
 charset EF1a_3 = 491-1100\3;
 charset RPB1_3_RPB2_3 = 2858-4619\3; charset INDEL = 1-329;
 partition favored = 7: ITS1_ITS2_I3_RPB1, 58S_EF1a_2_RPB1_2_RPB2_2, LSU, EF1a_1_RPB1_1_RPB2_1, EF1a_3, RPB1_3_RPB2_3, INDEL;
 set partition = favored;
 lset applyto=(1,3,4,5) nst=6 rates=invgamma ngammacat=4;
 lset applyto=(6) nst=6 rates=gamma ngammacat=4;
 lset applyto=(2) nst=2 rates=invgamma ngammacat=4;
 lset applyto=(7) rates=gamma;

 prset applyto=(1)
 statefreqpr=fixed(0.210937,0.227899,0.237667,0.323497)
 shapepr=fixed(0.873065)
 pinvar=fixed(0.218331)
 revmat=fixed(1.65262,5.79093,2.15199,0.799527,5.85249, 1);

 prset applyto=(2)
 statefreqpr=fixed(0.304123,0.220815,0.203096,0.271967)
 shapepr=fixed(0.649635)
 pinvar=fixed(0.709309)
 revmat=fixed(1, 2.66487, 1, 1, 2.66487, 1);

 prset applyto=(3)
 statefreqpr=fixed(0.263434,0.194104,0.297374,0.245088)
 shapepr=fixed(0.536896)
 pinvar=fixed(0.556043)
 revmat=fixed(1.49914,6.55845,2.31922,0.676263,14.4049, 1);

 prset applyto=(4)
 statefreqpr=fixed(0.261993,0.234213,0.352613,0.151181)
 shapepr=fixed(0.797382)
 pinvar=fixed(0.548203)
 revmat=fixed(1.13561,1.77856,0.720257,0.482009,6.51285, 1);

 prset applyto=(5)
 statefreqpr=fixed(0.124654,0.392391,0.247792,0.235163)
 shapepr=fixed(1.76944)
 pinvar=fixed(0.109357)
 revmat=fixed(2.75686,8.85939,0.0001,2.26517,21.744, 1);

 prset applyto=(6)
 statefreqpr=fixed(0.139993,0.3014,0.312622,0.245986)
 shapepr=fixed(1.40007)
 revmat=fixed(2.68707,18.3327,5.29837,1.80452,19.1293, 1);

 prset applyto=(7) ratepr=variable;

end;
~~~

**Suppdata1; Tree 1.**
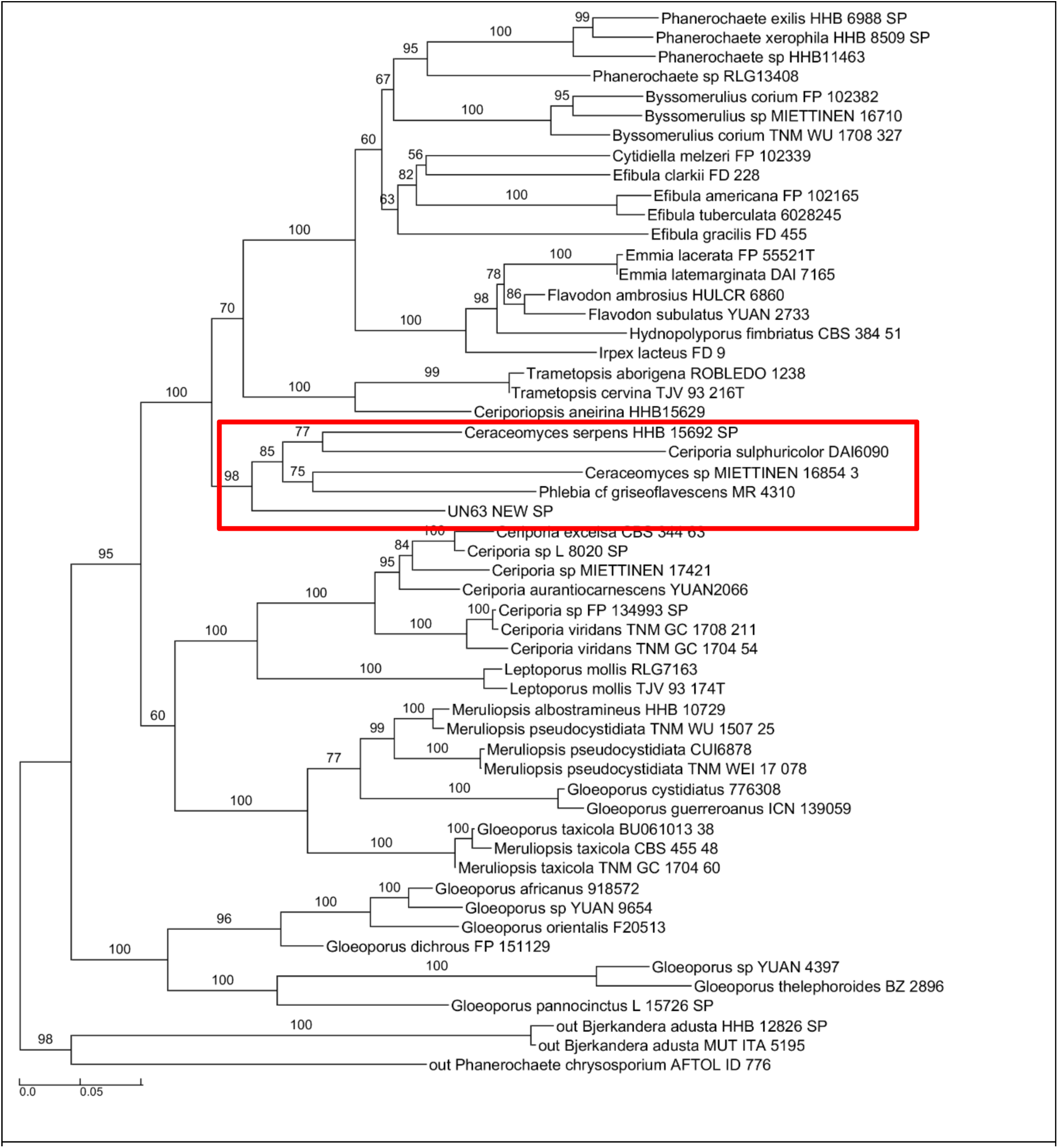
Maximum likelihood tree of Irpicaceae. Supporting values on branches are Ultrafast Bootstrap. Red box shows the new genus and species clade. Scale bar: nucleotide substitutions per site.

**Suppdata1; Tree 2.**
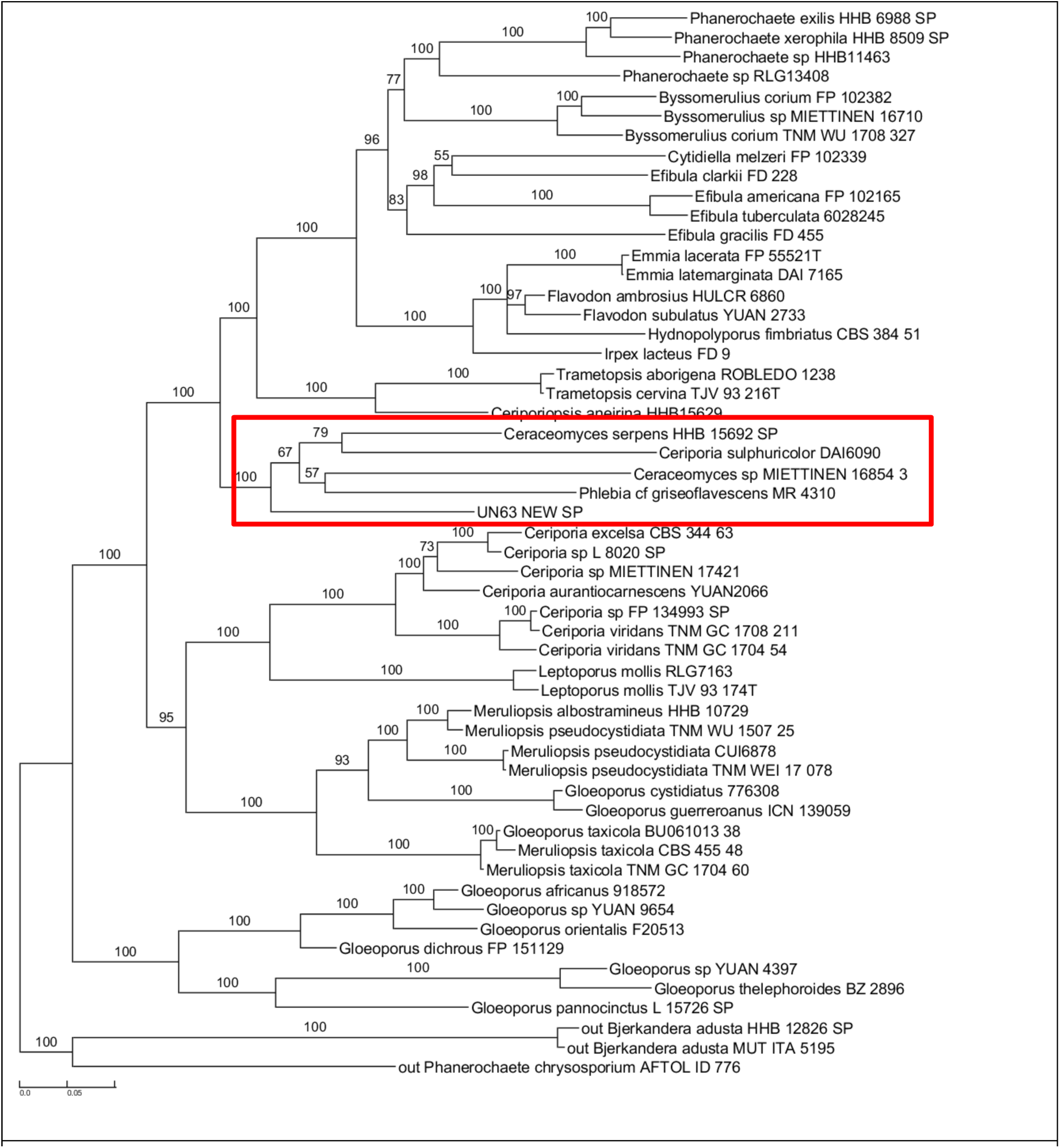
Bayesian Inference tree of Irpicaceae. Supporting values on branches are Bayesian Posterior Probability (in %). Red box shows the new genus and species clade. Scale bar: nucleotide substitutions per site.

**Suppdata1; Tree 3.**
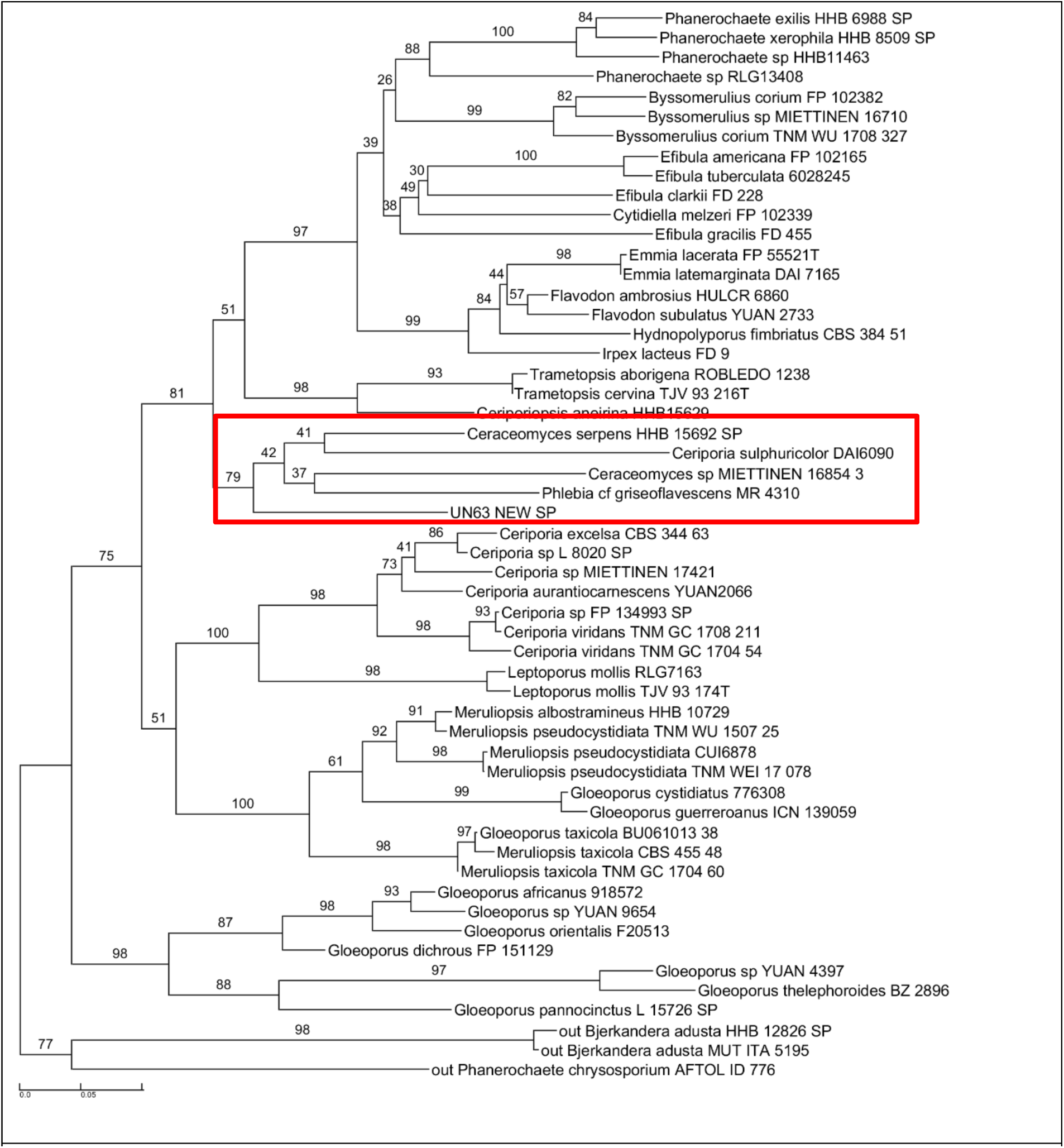
Maximum likelihood tree of Irpicaceae. Supporting values on branches are Non-parametric Bootstrap. Red box shows the new genus and species clade. Scale bar: nucleotide substitutions per site.

**Suppdata1; Tree 4.**
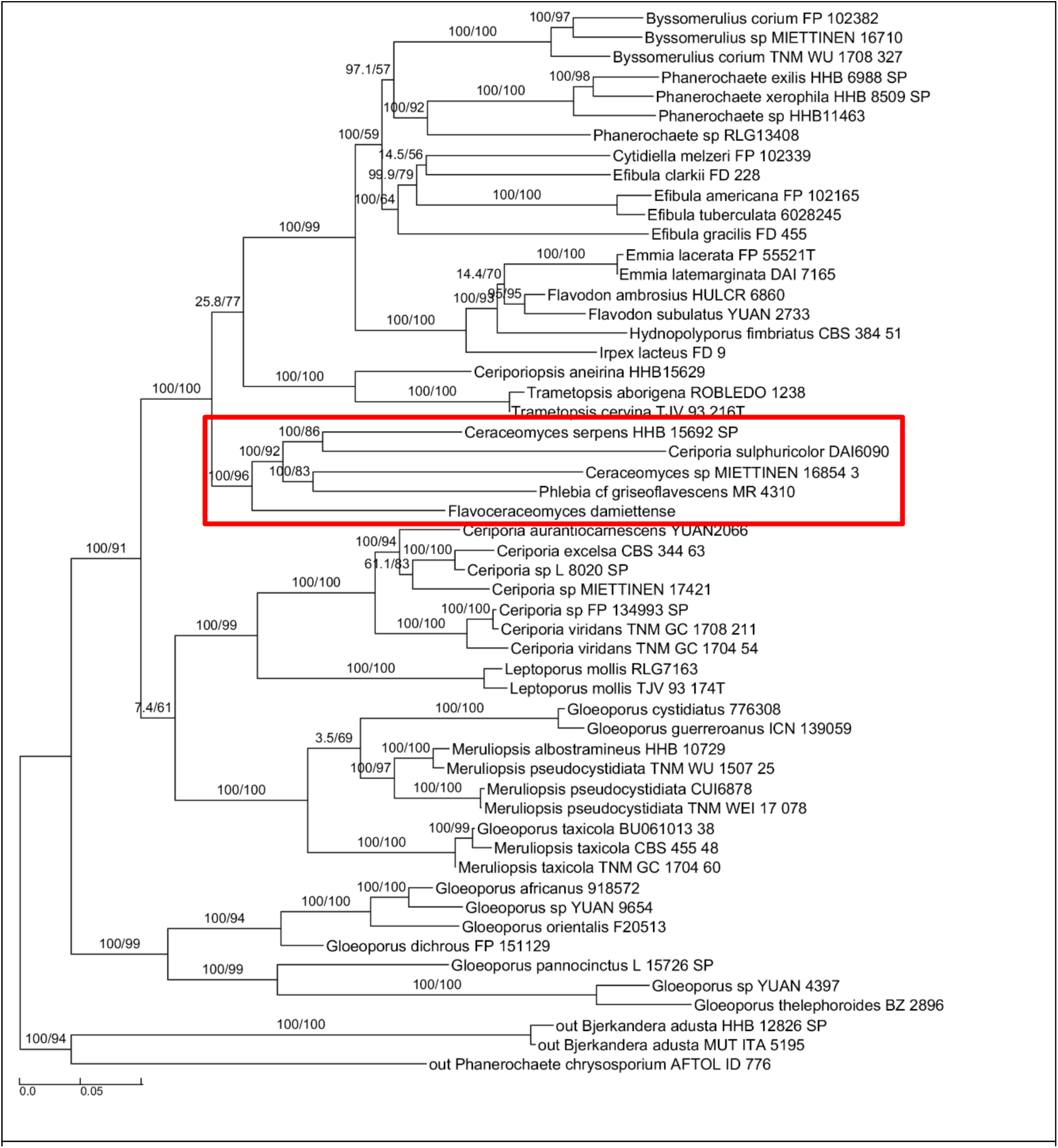
Maximum likelihood tree of Irpicaceae. Supporting values on branches are SH-alrt / Ultrafast Jackknife. Red box shows the new genus and species clade. Scale bar: nucleotide substitutions per site.

**Suppdata1; Tree 5.**
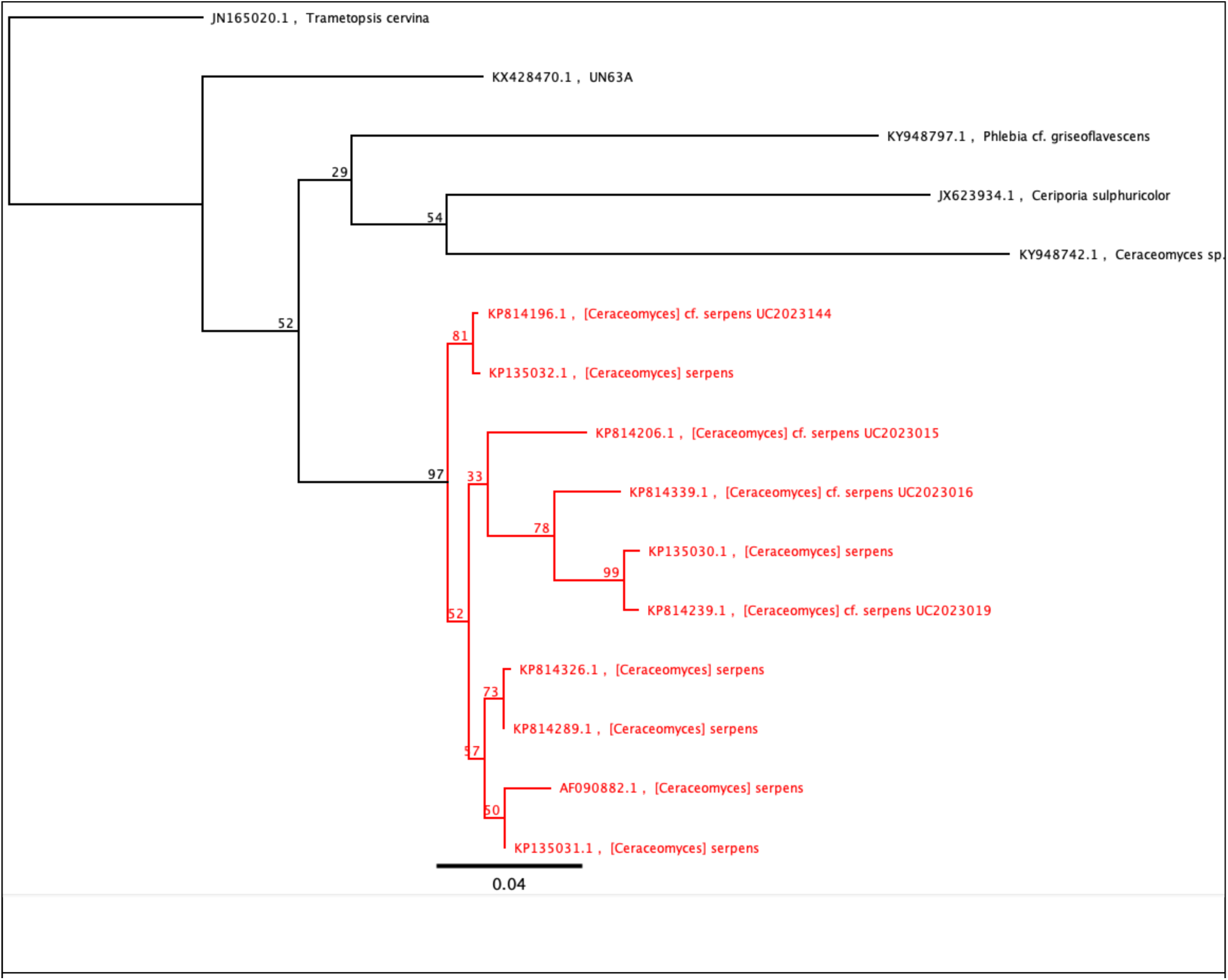
Maximum-likelihood tree of *Flavoceraceomyces* using ITS only including several sequences of *C. serpens*. This tree supports the position of *C. serpens* clustered within *Flavoceraceomyces*. Note that this tree was computed using Genious Prime.

## Notes

### Competing Interest Statement

The authors have declared no competing interest.

